# CryoTransformer: A Transformer Model for Picking Protein Particles from Cryo-EM Micrographs

**DOI:** 10.1101/2023.10.19.563155

**Authors:** Ashwin Dhakal, Rajan Gyawali, Liguo Wang, Jianlin Cheng

**Affiliations:** Department of Electrical Engineering and Computer Science, University of Missouri, Columbia, MO 65211, USA; NextGen Precision Health, University of Missouri, Columbia, Columbia, MO 65211, USA; Laboratory for BioMolecular Structure (LBMS), Brookhaven National Laboratory, Upton, NY 11973, USA

**Author notes:** Corresponding author: Jianlin Cheng.

## Abstract

Cryo-electron microscopy (cryo-EM) is a powerful technique for determining the structures of large protein complexes. Picking single protein particles from cryo-EM micrographs (images) is a crucial step in reconstructing protein structures from them. However, the widely used template-based particle picking process requires some manual particle picking and is labor-intensive and time-consuming. Though machine learning and artificial intelligence (AI) can potentially automate particle picking, the current AI methods pick particles with low precision or low recall. The erroneously picked particles can severely reduce the quality of reconstructed protein structures, especially for the micrographs with low signal-to-noise (SNR) ratios. To address these shortcomings, we devised CryoTransformer based on transformers, residual networks, and image processing techniques to accurately pick protein particles from cryo-EM micrographs. CryoTransformer was trained and tested on the largest labelled cryo-EM protein particle dataset - CryoPPP. It outperforms the current state-of-the-art machine learning methods of particle picking in terms of the resolution of 3D density maps reconstructed from the picked particles as well as F1-score and is poised to facilitate the automation of the cryo-EM protein particle picking.

## 1. Introduction

Cryogenic electron microscopy (cryo-EM) is a modern biophysical technique that captures two-dimensional (2D) images of biological macromolecules, such as proteins and viruses at cryogenic temperature [1], through the use of an electron detection camera. When subjected to an electron beam within a thin vitrified sample, this technique generates 2D image projections of the specimens (e.g., protein particles). These 2D representations are stored in various image formats (like mrc, tiff, tbz, eer, png, etc.), which are called micrographs. A single micrograph can contain hundreds or thousands of particles of a protein, randomly oriented in different directions. Given the inherent challenges of ascertaining the orientations of the particles and the low SNR of micrographs, hundreds of thousands of high-quality particles are often required to be identified to determine a high-resolution three-dimensional (3D) structure of the protein.

The initial step of determining the 3D structure of the proteins from the micrographs involves the recognition and extraction of particles from 2D micrographs, which is commonly referred to as particle picking. Its primary goal is to identify and locate individual protein particles within each micrograph while excluding malformed particles, crystalline ice contamination, and background regions. Essentially, the task of particle picking involves taking a micrograph as input and generating the coordinates for all protein particles present in that micrograph as the desired output (stored in the form of .*box* or .*star* files). These coordinates thereafter serve as the data for subsequent stages of 3D protein structure reconstruction. These 3D structures of proteins are important for understanding their biological functions [2] and their interactions with ligands [3], [4], facilitate structure-based drug discovery [3] [5].

Because the SNR of micrographs is generally low, hundreds to thousands of micrographs need to be generated to obtain a high-resolution structure for a protein, from which as many as millions of particle images can be picked. Precise identification of true particles is important, as the presence of false positive particles complicates the down-stream 3D protein reconstruction process. The particle picking task is inherently challenging due to several factors, including high noise levels caused by ice and contamination, low contrast of particle images, heterogenous conformations of particles, and variation in the orientation of particles.

This manual picking process by human is laborious, tedious, and time-consuming, which cannot be applied to pick millions of particles from thousands of micrographs. Therefore, substantial efforts have been put to develop semi-automated or fully automated methods to pick protein particles, which can be classified into two categories: (1) template-based particle picking and (2) machine learning particle picking.

In the template-based particle picking, the identification of particles primarily hinges on measuring a potential particle’s similarity to user-predefined (manually selected) reference particles called templates. Because micrographs are usually noisy due to various factors such as ice contamination, carbon areas, overlapping particles, and other impurities, the template-based particle picking is often unable to detect particles of unusual shape and suffers from high false-positive rates. As a result, subsequent steps of manual particle selection are necessary to filter the particles picked by the template-based particle picking. Typically, iterative 2D-3D classification techniques are employed to scrutinize the picked particles and remove false particles. However, this particle picking, and downstream manual curation may introduce a degree of human bias into the final particle set selection, which may mistakenly exclude rare particle views and distinct conformations that are important for building high-resolution protein structures. Thus, this approach generally necessitates a large degree of human intervention and trial and errors to obtain good results.

The machine learning particle picking consists of both unsupervised learning (clustering) methods [6] and supervised learning methods [7] [8] [9] [10]. Recently, a number of deep learning methods were developed to automate the protein particle picking, which include XMIPP [11], DeepPicker [12], DeepEM [13], Xiao et al.’s method [14], Warp [15], HydraPicker [16], McSweeney et al.’s method [17], DRPnet [18], CrYOLO [19] and Topaz [20]. Among them, CrYOLO and Topaz based on convolutional neural networks have been widely used in particle picking. However, they have been trained with limited particle data and have the difficulty to generalize to new protein types or shapes. For instance, CrYOLO usually overlooks many true protein particles, while Topaz often picks excessive numbers of duplicate particles and some false positives such as ice contaminants and false particles in carbon-rich areas.

To overcome these obstacles, we devised a transformer-based particle picking approach and trained it on the largest, diverse, manually-labelled CryoPPP protein particle dataset [21], [22]. Inspired by Meta’s Detection Transformer (DETR) [23] for detecting small objects, we designed the end-to-end detection transformer named as CryoTransformer. Briefly, it has an initial step of reducing noise in micrographs (**Figure 1A, B**), followed by the feature extraction through a ResNet-152 architecture (**Figure 1C**). Subsequently, a transformer model is used for detecting protein particles as shown in **Figure 1D**. This is succeeded by the feed-forward networks to predict particles (**Figure 1E**), which are followed by the post-processing procedures. The output (**Figure 1F**) includes particle markings on the micrographs stored in.*star* files, which can be directly used for the subsequent stages of 3D protein structure reconstruction. We conducted a rigorous evaluation of CryoTransformer. It outperforms the two popular deep learning methods: CrYOLO and Topaz. The source code and data for CryoTransformer are openly available at: https://github.com/jianlin-cheng/CryoTransformer.

**Figure 1:**
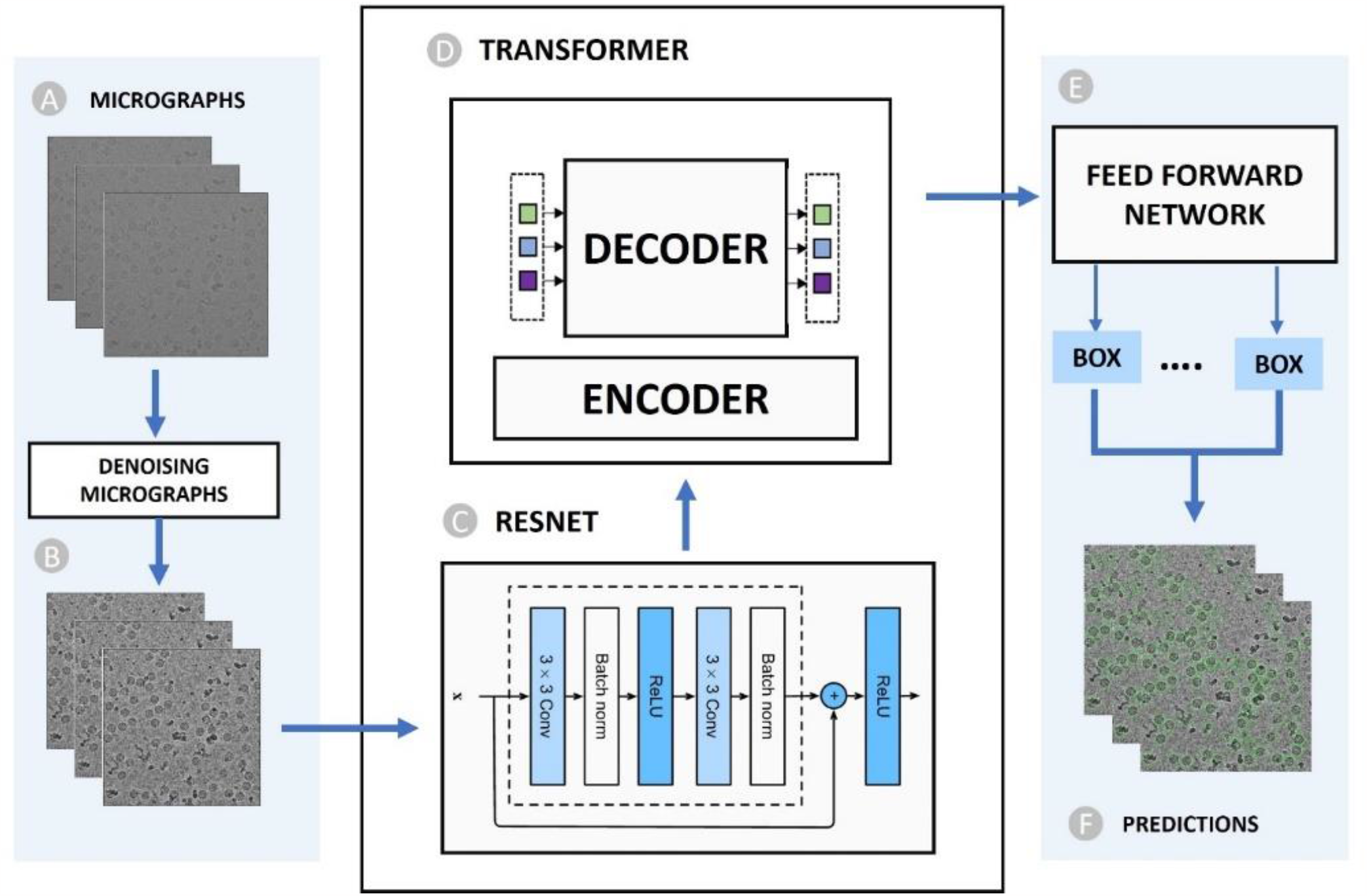
Overview of the CryoTransformer Particle Picking Pipeline. (A) Input raw micrograph undergoes initial denoising. (B) Denoised micrographs serve as input for subsequent processing. (C) CNN-Based Resnet-152 architecture extracts image features. Features extracted in (C) are processed by an (D) encoder-decoder Transformer. (E) Feed-forward networks further refine the processed data. (F) Predictions of particles encircled in micrographs, eventually stored in star files as the final output.

## 2. Materials and Methods

### 2.1 Dataset

### Dataset acquisition

We utilized the largest comprehensive CryoPPP dataset [21], [22] curated from Electron Microscopy Public Image Archive (EMPIAR) [24], to train, validate, and test CryoTransformer. The micrographs of 22 proteins (EMPIAR IDs) from the CryoPPP dataset were used, with the data of each EMPIAR ID split according to an 80%-10%-10% ratio for training, validation, and internal test. Moreover, we used the data of 6 distinct EMPIAR IDs in CryoPPP dataset different from the 22 proteins above as well as the 4 complete micrograph datasets from EMPIAR repository [24] as the independent test dataset to compare CryoTransformer with the external methods.

The selection of training and test data considered a range of protein attributes, including type, shape, size, and overall structural characteristics. The 22 proteins used for the training, validation and internal test are described in **Table 1. Supplementary Figure S1** illustrates the varying defocus values of the training data. The datasets encompass various protein categories, such as transport proteins, membrane proteins, viral proteins, ribosomes, signaling proteins, aldolases, and more. They are comprised of micrographs featuring diverse attributes, including those with ice patches, contaminants, varying ice thickness, and carbon areas. Different protein distribution patterns, including monodisperse, clumped clusters, and heterogeneous views, are also included. The **Supplementary Table S1 and S2** contain the information and statistics of the proteins in the independent test dataset.

**Table 1:** The statistics and information of the 22 sets of micrographs for training, validation, and internal test of CryoTransformer (* Theoretical weight)

| SN | EMPIAR ID | Type of Protein | Image Size | Total Structure Weight (kDa) | # Training Micrographs | # Validation Micrographs | # Test Micrographs | # Total Micrographs |
| --- | --- | --- | --- | --- | --- | --- | --- | --- |
| 1 | 11183 [25] | Signaling Protein | (5760, 4092) | 139.36 | 250 | 25 | 25 | 300 |
| 2 | 11057 [26] | Hydrolase | (5760, 4092) | 149.43 | 250 | 25 | 20 | 295 |
| 3 | 11051 [27] | Transcription/DNA/RNA | (3838, 3710) | 357.31 | 250 | 25 | 25 | 300 |
| 4 | 10852 [28] | Signaling Protein | (5760, 4092) | 157.81 | 270 | 40 | 33 | 343 |
| 5 | 10816 [29] | Transport Protein | (7676, 7420) | 166.62 | 250 | 25 | 25 | 300 |
| 6 | 10760 [30] | Membrane Protein | (3838, 3710) | 321.69 | 250 | 25 | 25 | 300 |
| 7 | 10737 [31] | Membrane Protein | (5760, 4092) | 155.83 | 250 | 25 | 17 | 292 |
| 8 | 10671 [32] | Signaling Protein | (5760, 4092) | 77.14 | 250 | 25 | 23 | 298 |
| 9 | 10590 [33] | Transport Protein | (3710, 3838) | 1000* | 250 | 25 | 21 | 296 |
| 10 | 10526 [34] | Ribosome (50S) | (7676, 7420) | 1085.81 | 180 | 20 | 20 | 220 |
| 11 | 10444 [35] | Membrane Protein | (5760, 4092) | 295.89 | 250 | 25 | 21 | 296 |
| 12 | 10406 [36] | Ribosome (70S) | (3838, 3710) | 632.89 | 200 | 20 | 19 | 239 |
| 13 | 10387 [37] | Viral Protein | (3710, 3838) | 185.87 | 250 | 25 | 24 | 299 |
| 14 | 10291 [38] | Transport Protein | (3710, 3838) | 361.39 | 250 | 25 | 25 | 300 |
| 15 | 10289 [38] | Transport Protein | (3710, 3838) | 361.39 | 250 | 25 | 25 | 300 |
| 16 | 10240 [39] | Lipid Transport Protein | (3838, 3710) | 171.72 | 250 | 25 | 24 | 299 |
| 17 | 10184 [40] | Aldolase | (3838, 3710) | 150* | 250 | 25 | 21 | 296 |
| 18 | 10096 [41] | Viral Protein | (3838, 3710) | 150* | 250 | 25 | 25 | 300 |
| 19 | 10077 [42] | Ribosome (70S) | (4096, 4096) | 2198.78 | 250 | 25 | 25 | 300 |
| 20 | 10075 [43] | Bacteriophage MS2 | (4096, 4096) | 1000* | 250 | 25 | 24 | 299 |
| 21 | 10059 [44] | Transport Protein | (3838, 3710) | 317.88 | 250 | 25 | 16 | 291 |
| 22 | 10005 [45] | Transport Protein | (3710, 3710) | 272.97 | 22 | 4 | 3 | 29 |
|  |  | **Total Micrographs** |  |  | **5,172** | **534** | **486** | **6,192** |

### Denoising and pre-processing of cryo-EM micrographs

The cryo-EM micrographs in .*mrc* format, serve as the initial input for CryoTransformer. To reduce noise and improve the signal-to-noise ratio, a Gaussian filter with a kernel size of 9 is applied to convolve with the images. Subsequently, the images undergo standard normalization to achieve consistent intensity ranges. The normalized pixel values of the images are computed using the formula [pixel = (pixel-μ)/ σ], ensuring that the data is centered and scaled appropriately for the further analysis. The normalized images are then converted to grayscale, which collapses multi-channel intensity information into a single channel, ensuring a uniform representation of pixel values ranging from 0 to 255 (**Figure 2A**).

Effective noise reduction is essential to reveal clear structural details in cryo-EM micrographs. We employ a two-step denoising process to the normalized images, involving Fast Non-Local Means (FastNLMeans) denoising followed by Weiner filtering (**Figure 2B**). FastNLMeans denoising is employed to retain image details while suppressing noise artifacts. By exploiting the redundancy present in natural images, FastNLMeans replaces the noisy pixel with a weighted average of similar pixels from a larger neighborhood. The trade-off between noise suppression and detail preservation is controlled by the choice of template window size (7 in this case) and the search window size (21 in this case).

**Figure 2:**
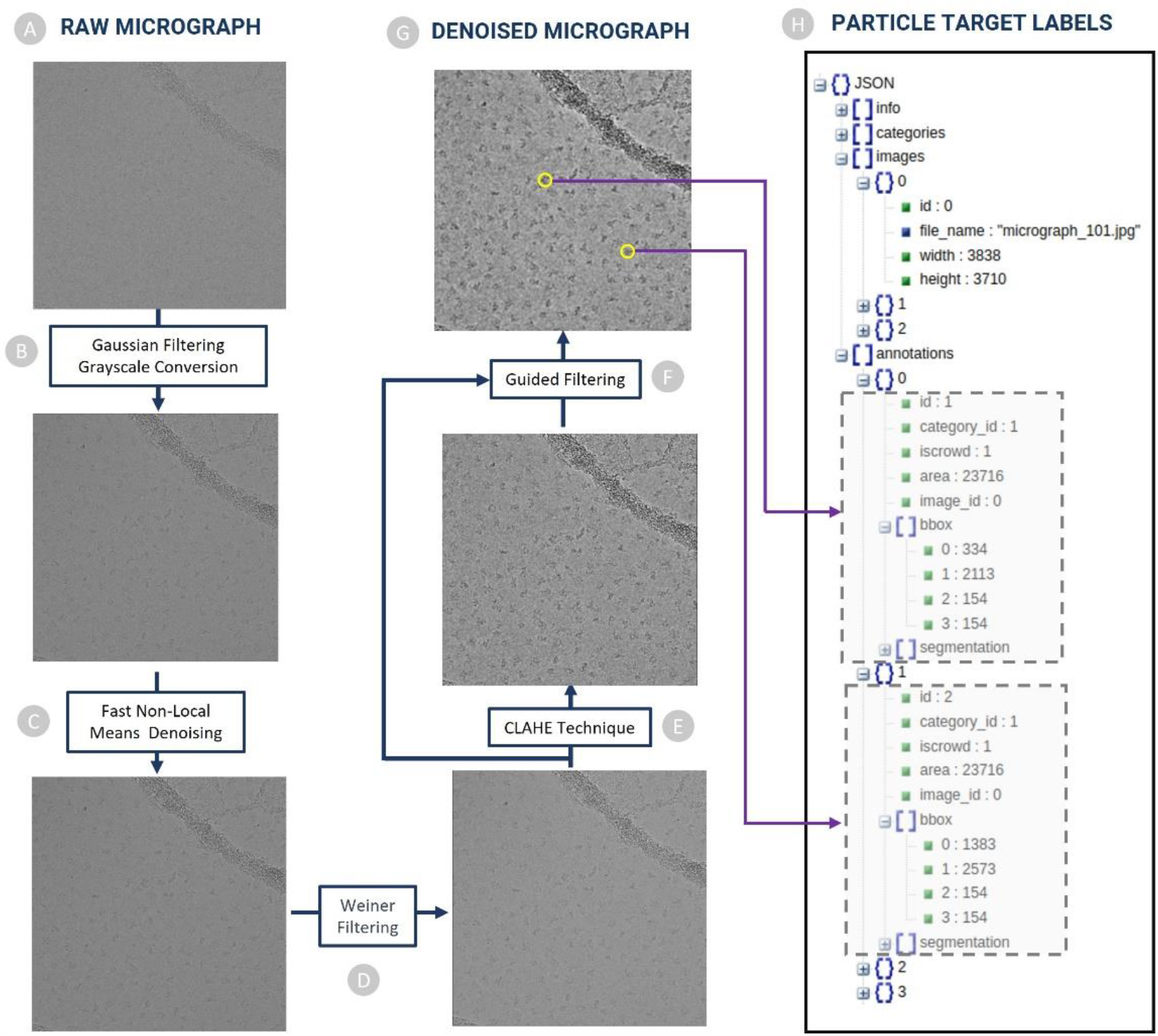
The denoising used to preprocess cryo-EM micrographs in CryoTransformer. (A) Raw micrographs with low contrast and low SNR go through (B) Gaussian filtering and Grayscale conversion. Normalized micrograph undergoes (C) FastNLMeans denoising technique. (D) Weiner filtering is applied to the micrographs from the previous step, and subsequently (E) CLAHE technique is used to enhance visual clarity of the micrograph. Eventually, (F) Guided filtering is performed using the CLAHE-enhanced image as a guide to obtain (G) denoised micrographs. (H) Ground truth particle annotation data. Particle coordinates from ground truth coordinate files are extracted to create COCO-dataset that is used as target labels for training CryoTransformer.

The output of FastNLMeans denoising is subjected to Weiner filtering to further reduce the residual noise and enhance the image’s structural fidelity (**Figure 2C**). It achieves this by estimating the original image’s frequency spectrum and applying a correction factor to mitigate the effects of noise. Enhancing contrast in cryo-EM micrographs is crucial for improving particle visibility and overall image quality. We incorporate the Contrast Limited Adaptive Histogram Equalization (CLAHE) technique for this purpose (**Figure 2D**). The CLAHE technique, with a clip limit of 2 and a tile grid size of 16x16, is applied to the denoised images. This technique effectively addresses non-uniform illumination and low contrast, leading to enhanced visual clarity.

To accomplish selective smoothing and fine detail preservation, guided filtering is performed using the CLAHE-enhanced image as a guide (**Figure 2E**). Guided filtering operates by estimating the local linear relationship between the guidance image and the target image (**Figure 2F**). This relationship is then used to determine the filtering weights applied to each pixel, resulting in controlled smoothing, while retaining sharp edges and fine details. The filtering fine-tunes the micrographs, achieving a balance between noise reduction and preservation of important structural information (**Figure 2G**).

### senerating COCO-dataset for labelled protein particles in micrographs

We used the ground truth particle coordinate data from the CryoPPP dataset [21], [22] to generate labels to train CryoTransformer. The particle labels were stored in the widely adopted Common Objects in Context (COCO) format [46]. This format is extensively used for object detection and segmentation tasks, and it adheres to a structured JSON layout that defines how labels and associated metadata are stored for an image dataset. An illustration of how these labels are stored is depicted in **Figure 2H**. In the case of all training and validation images, we have two JSON files: one for training (referred to as the “train JSON”) and another for validation (referred to as the “validation JSON”). We chose to adopt this labeling data format because the COCO format imposes a standardized structure for annotations, including object category labels and bounding box coordinates. This uniformity streamlines the data preprocessing process and ensures that models can readily comprehend and learn from the annotated data. The COCO format permits the annotation of multiple objects (protein particles) within a single image (micrograph). Each object is associated with its distinct category label and bounding box. For each particle, we retain details such as its bounding box coordinates, area, category label (typically set to 1 in our case as all objects to be detected are protein particles), the corresponding image reference, and a unique particle ID.

### 2.2 Design and Implementation of CryoTransformer

CryoTransformer is designed to achieve the accurate prediction of bounding boxes for the protein particles within a micrograph, while minimizing the number of false positives. It undergoes an end-to-end training, using a specialized loss function that effectively combines the bipartite matching loss between predicted and ground-truth protein particles in the micrographs.

### CryoTransformer Architecture

As illustrated in **Figure 3**, CryoTransformer comprises three main components: a Convolutional Neural Network (CNN) with residual connections (Resnet-152 [47]) responsible for feature extraction, an encoder-decoder transformer [23], [48] for learning the shapes of the particles in the context of an entire image, and a feed-forward network (FFN) responsible for producing the ultimate particle predictions.

**Figure 3:**
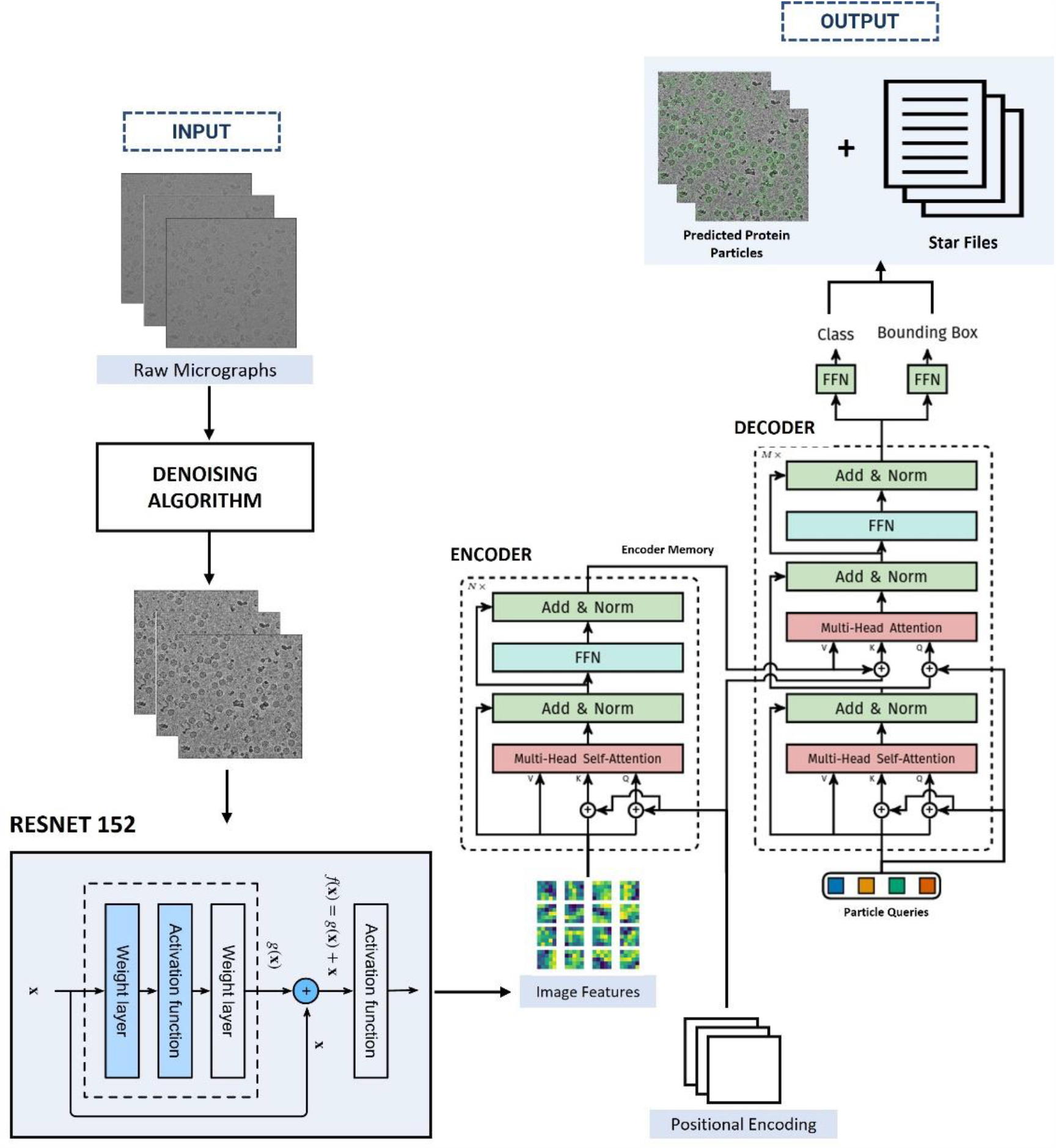
Architecture of CryoTransformer. The raw micrographs are denoised and are fed into the ResNet-152 module for feature extraction. The images features, along with positional encoding, are fed to the encoder of the transformer. The output from the encoder is subsequently passed to the decoder layer. Finally, the decoder’s output is passed to the feed forward networks that generate the protein particle bounding box predictions. These predictions are used in generating the predicted protein particles encircled in micrographs, which are stored in the .star files.

### Resnet-152 Backbone Block

The Resnet-152 receives the preprocessed micrographs 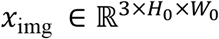 (with 3 color channels)) as input and generates a lower-resolution activation map as *f* ∈ ℝ^*C*×*H*×*W*^, Where *C* = 2048, and 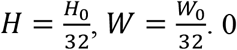 0 padding is applied to the images in a batch to make sure that they all have same input dimensions (*H*_0_, *W*_0_) as the largest image size of the batch.

### Transformer Module

The features extracted from the Resnet-152 are subsequently passed through the transformer. This transformer consists of two main components: encoder and decoder. The image features from the Resnet-152 backbone block are passed through the transformer along with the positional encoding and particle queries. The transformer outputs intermediate predictions, which are fed to the FFN module to predict particle labels and bounding boxes.

#### Transformer Encoder

The encoder plays a vital role in generating coherent and context-aware outputs. In the encoder, a 1x1 convolution operation is used to decrease the channel dimension of the high-level activation map, denoted as *f*, from *C* to a smaller dimension *d*, yielding a new feature map *z*_0_ ∈ ℝ^*d*×*H*×*W*^. Since the encoder accepts a one-dimensional sequence as input, we collapse the spatial dimensions of *z*_0_ into a single dimension. As a result, the resultant input becomes a feature map of dimension *d* × *HW*. Here, every encoder layer follows a consistent structure, comprising a multi-head self-attention component and a FFN layer. To account for the permutation-invariant nature of the transformer architecture, we enhance it by incorporating the positional encodings [49] [50], which are included in the input of every multi-head self-attention layer.

#### Transformer Decoder

The decoder receives the memory from encoder, positional encoding, and particle queries as input. It involves the transformation of *N* embeddings of size d (in our specific scenario, *N* = 600, meaning predicting max 600 protein particles per micrograph) through the multi-headed self-attention mechanisms. It’s worth noting that since the decoder is also designed to be permutation-invariant, it requires distinct particle queries (initialized as random vectors) within the set of *N* inputs to generate different outcomes. These particle queries, added to the input at each attention layer, are a are updated through back propagation. Subsequently, the output of the decoder is individually used to predict box coordinates and class labels (1 in our case) through a feed-forward network, a process detailed in the following subsection, resulting in *N* final predictions.

### Feed-Forward Networks Module

The final prediction is generated through a 3-layer perceptron with a ReLU activation function and *d* hidden nodes in each hidden layer, followed by a linear projection layer. This FFN is responsible for predicting the normalized center coordinates, height, and width of the bounding box relative to the input micrograph. Additionally, the linear layer predicts the class label using a softmax function. Considering that we are making predictions for a fixed-size set of *N* potential bounding boxes, and *N* is typically much larger than the actual number of protein particles in a single micrograph, we introduce a special class label denoted as *∅*. This label means that no protein particle has been detected in a particular slot. Its role is akin to the “background” class in conventional object detection.

### Loss Function

CryoTransformer generates a consistent set of *N* predictions in a single traversal of the decoder. This number *N* was deliberately chosen to exceed the usual count of protein particles in a micrograph. To achieve this, the loss function is designed to establish an ideal bipartite matching between the predicted protein particles and their corresponding ground truth. Subsequently, the model optimizes the losses pertaining to individual particles in order to refine the predictions further.

We can represent the ground truth set of particles as *y* and the set of *N* predictions as 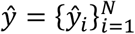. When N exceeds the number of true protein particles in the micrograph, we enlarge *y* as a set of size *N*, with padding represented by *∅* (no protein particle). To find the optimal bipartite matching between these two sets, we aim to find a permutation of *N* elements denoted as *σ* ∈ *𝔖*_N_ that incurs the lowest cost. This permutation is determined by the following equation, given in equation I:

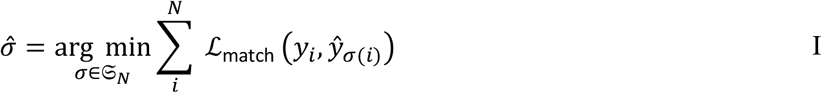

ℒ_match_ (*y*_*i*_, *ŷ*_*σ*_(*i*)) represents the pairwise matching cost between the ground truth particle *y*_*i*_ and a prediction indexed by *σ*(*i*). This cost is calculated using the following equation II:

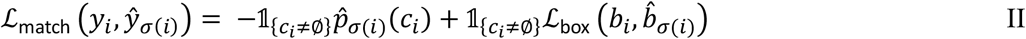

We can view each element *i* in the ground truth set as a *y*_*i*_ = (*c*_*i*_, *b*_*i*_), where *c*_*i*_ represents the target class label, and *b*_*i*_ belongs to the range [0,1]^4^, representing a vector that specifies the center coordinates of the ground truth box, along with its height and width relative to the micrograph dimensions. This approach ensures a one-to-one matching, preventing duplicate predictions when directly predicting sets.

The next stage involves calculating the Hungarian loss using the Hungarian algorithm [51] for all pairs that were matched in the preceding step. We define this loss according to the equation III:

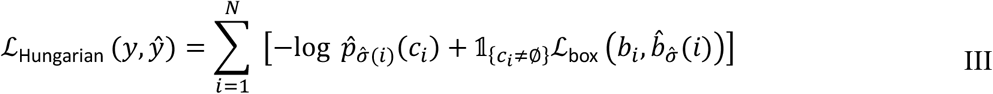

Here, 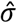 represents the optimal assignment obtained from the initial equation I.

In practical implementation, we apply a down-weighting factor of 10 to the log-probability term when *c*_*i*_ is equal to *∅*, denoting the absence of a particle. This adjustment is made to address the issue of class imbalance. The second part of the Hungarian loss (ℒ_box_ (⋅)) scores the bounding boxes is given by the equation IV:

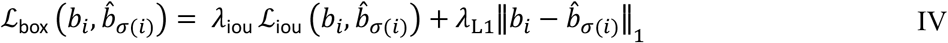

Where *λ*_iou_, *λ*_L1_ ∈ ℝ are hyperparameters and ℒ_iou_ (⋅) is the generalized IoU [52] given by equation V:

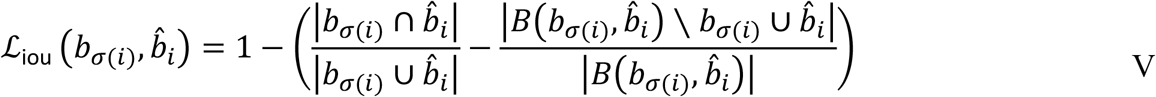

In the context provided, |.| denotes “area,” and we use the terms union and intersection of box coordinates as shorthand references for the boxes themselves. To compute the areas of unions or intersections, we rely on the minimum/maximum of linear functions involving *b*_σ_(*i*)and 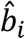. This approach ensures that the loss behaves in a stable manner for the computation of stochastic gradients. 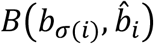 refers to the largest bounding box that contains both 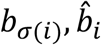.

### Model Implementation and Training

We trained CryoTransformer with AdamW optimizer [52] by setting the initial transformer’s learning rate to 10^−4^, the backbone’s to 10^−5^, and weight decay to 10^−4^. All weights are randomly initialized with Xavier initialization [53]. Additive dropout of 0.1 is applied after every multi-head attention and FFN before layer normalization. We use a training schedule of 300 epochs with a learning rate drop by a factor of 10 after 200 epochs, where a single epoch is a pass over all training images once. Training the model for 300 epochs on NVIDIA A100 80GB GPU took 2 days and 11 hours to complete.

### 2.3 Postprocessing Predictions and Reconstructing Protein Density Maps from Picked Particles

The FFN module of CryoTransformer predicts the coordinates of particles and their corresponding confidence scores (ranging from 0 to1). The predictions are processed in a few steps to generate final particle predictions, group the picked particles into different 2D orientation classes, and use them to build 3D density maps of proteins. The visual representation of the overall process is shown in **Figure 4**.

**Figure 4:**
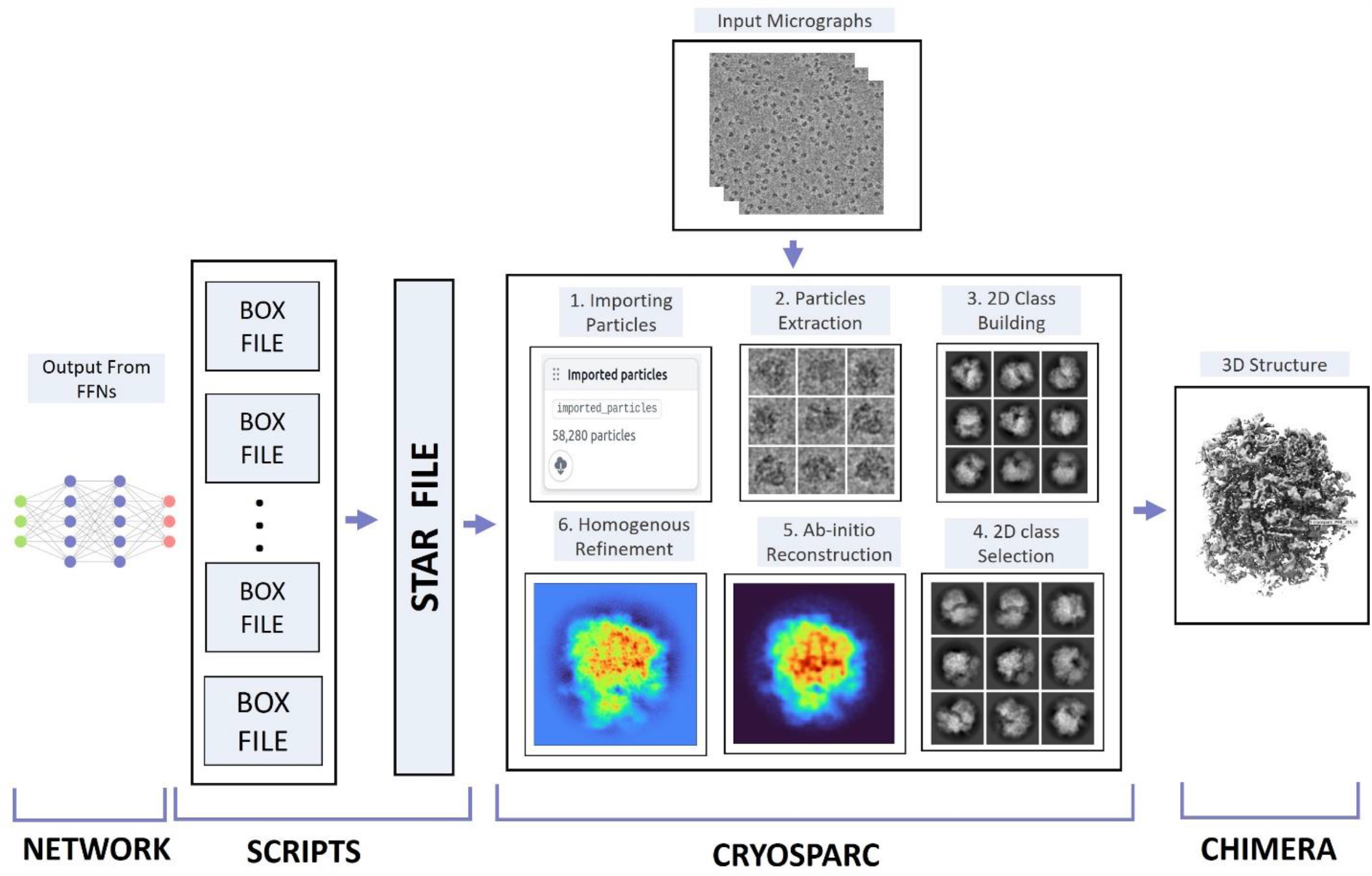
Post processing steps to generate 3D protein density maps from picked particles. CryoTransformer outputs the .star files, which are imported into CryoSPARC along with the micrographs. Steps (1-6) are performed in CryoSPARC to generate the 3D density maps for a protein. The resolution of the density maps is employed as the main metric to evaluate the quality of the picked particles.

The predictions are first used to generate individual box files for every micrograph for a protein, containing the center coordinates (x and y) of all the predicted protein particles. We retain only the particles whose confidence score falls in the range from 25^th^ percentile to 100^th^ percentile. Subsequently, these box files are merged to create a .*star* file that can be accepted by CryoSPARC [54] for density map construction for the protein.

The star files generated are imported into CryoSPARC through the *‘import particles’* task, accompanied by input parameters such as Acceleration Voltage (kV), Spherical Aberration (mm), and Pixel Size (Å) as well as the patch-based Contrast Transfer Function (CTF)-estimated micrographs. Subsequently, these particles are extracted using a specified extraction box size (in pixels) and fed into the 2D classification function of CryoSPARC to group them into different orientation classes.

This 2D classification step helps identify and exclude false particles through manual inspection, which usually can improve the resolution of the density maps reconstructed from the picked particles.

To assess the quality of the particles picked by CryoTransformer, CrYOLO and Topaz, we carried out the density map reconstruction experiments with and without the 2D selection respectively. When the 2D classification was used, we generated a total of 50 particle classes, employing a window inner radius of 0.85 and an outer radius of 0.99. Additionally, we performed 15 iterations to refine the CryoSPARC’s noise model. The selected particles were used by an *ab initio* reconstruction process with the standard parameter settings, which includes 300 iterations of reconstruction with a Fourier radius step of 0.04 and a momentum of 0 and an initial learning rate of 0.4 for the stochastic gradient descent optimization. Additionally, a lowpass filter cutoff in Fourier radii of 7 was applied to the initial random structures.

After generating the initial density map for a protein, the cryoSPARC’s ‘*homogeneous refinement’* job was employed to enhance it further. The homogeneous refinement was applied to correct the higher-order aberrations and to refine particle defocus caused by factors such as beam tilt and spherical aberration. To ensure the fairness in comparisons of the particle picking methods, the experiment was conducted three times for each method with different random seed values, and the best score (in Angstrom units) out of the three experiments was used in the comparison.

## 3. Results

We evaluated the particle picking performance of CryoTransformer in the following complementary ways. First, we compared it with CrYOLO and Topaz in terms of the resolution of the density maps reconstructed from the particles picked by them from the full set of micrographs in the EMPIAR repository for the four proteins in the independent test dataset. Second, we compared it with CrYOLO and Topaz in terms of the resolution of the density maps picked from a subset of labeled micrographs in the CryoPPP dataset for the proteins in the independent test dataset. Finally, we visually inspected and assessed the particles picked by the three methods.

### 3.1 Comparing CryoTransformer, CrYOLO, and Topaz in terms of resolution of density maps reconstructed from the particles picked from the full set of micrographs in the EMPIAR repository (~1600 micrographs per protein)

The full set of micrographs in the EMPIAR repository for the four test proteins (Human HCN1 Hyperpolarization-Activated Channel (EMPIAR 10081), Influenza Hemagglutinin (EMPIAR 10532), mechanotransduction channel NOMPC (EMPIAR 10093), and asymmetric αVβ8 (EMPIAR 10345)) in the independent test dataset were used to compare CryoTransformer, CrYOLO and Topaz. The resolution of the density map reconstructed from the particles picked by each method for each protein was calculated. The density maps were reconstructed by CryoSPARC in two modes: with 2D particle selection (*Select 2D*) or without it. The experiment for each method and each protein was conducted three times and the best results were selected for the comparison. The comparative results of the three methods are summarized in **Table 2**, while the detailed results of each trial reported in **Supplementary Table S3**.

**Table 2:**
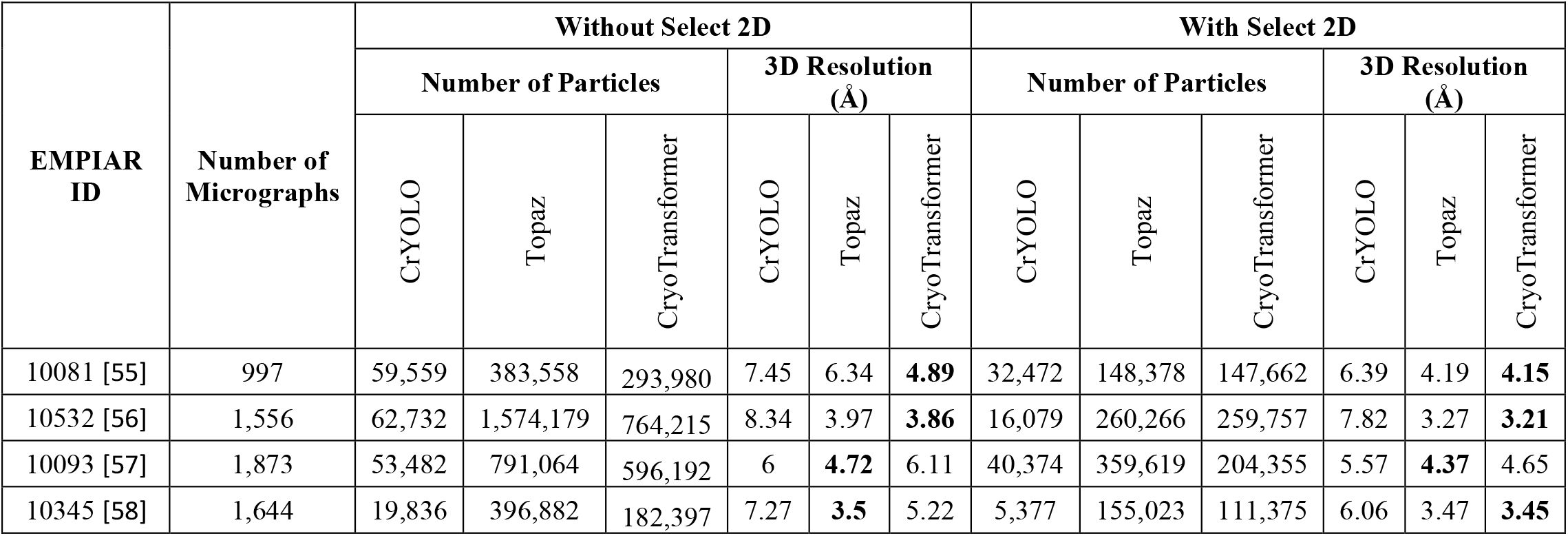
Comparison of CryoTransformer with crYOLO and Topaz’s performance in terms of the resolution of density maps reconstructed from the particles picked from the full set of micrographs of the four test proteins.

| EMPIAR ID | Number of Micrographs | Without Select 2D |  |  |  |  |  | With Select 2D |  |  |  |  |  |
| --- | --- | --- | --- | --- | --- | --- | --- | --- | --- | --- | --- | --- | --- |
|  |  | Number of Particles |  |  | 3D Resolution (Å) |  |  | Number of Particles |  |  | 3D Resolution (Å) |  |  |
|  |  | CrYOLO | Topaz | CryoTransformer | CrYOLO | Topaz | CryoTransformer | CrYOLO | Topaz | CryoTransformer | CrYOLO | Topaz | CryoTransformer |
| 10081 [55] | 997 | 59,559 | 383,558 | 293,980 | 7.45 | 6.34 | **4.89** | 32,472 | 148,378 | 147,662 | 6.39 | 4.19 | **4.15** |
| 10532 [56] | 1,556 | 62,732 | 1,574,179 | 764,215 | 8.34 | 3.97 | **3.86** | 16,079 | 260,266 | 259,757 | 7.82 | 3.27 | **3.21** |
| 10093 [57] | 1,873 | 53,482 | 791,064 | 596,192 | 6 | **4.72** | 6.11 | 40,374 | 359,619 | 204,355 | 5.57 | **4.37** | 4.65 |
| 10345 [58] | 1,644 | 19,836 | 396,882 | 182,397 | 7.27 | **3.5** | 5.22 | 5,377 | 155,023 | 111,375 | 6.06 | 3.47 | **3.45** |

With *Select 2D*, CryoTransformer has the highest resolution of the reconstructed density maps for three out of four proteins (i.e., EMPIAR IDs: 10081, 10532, and 10345), while Topaz has the highest resolution for one protein. Without *Select 2D*, CryoTransformer and Topaz each perform best on two proteins. The detailed assessment of crYOLO, Topaz, and CryoTransformer based on the 3D resolution of Gold Standard Fourier Shell Correlation (CSFSC) curves, 3D density maps, and density projections with *Select 2D* is visualized in **Figure 5**.

**Figure 5:**
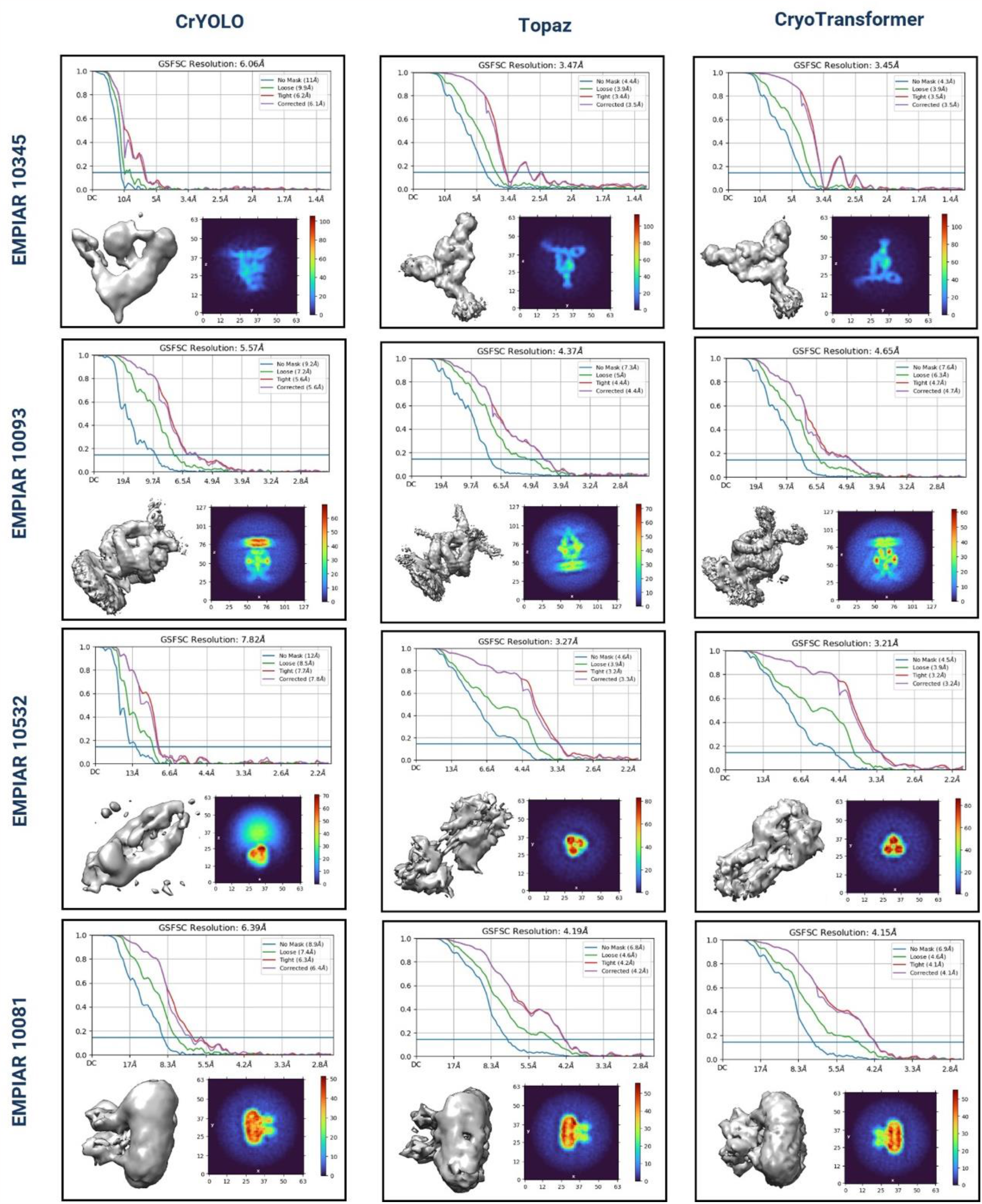
Assessment of CrYOLO, Topaz, and CryoTransformer based on the 3D resolution CSFSC curves, 3D density maps, and density projections. The top diagram in each row shows CSFSC curves, which indicate the resolution of 3D density maps for proteins structures reconstructed from picked particles. Bottom-left image in each sub-figure provides a visual representation of the 3D density map. The bottom-right image in each sub-figure depicts the density projections from the intermediate output of the ab initio reconstruction phase. The integrated density values along the normal direction to that plane are displayed. The color scheme in the heatmap corresponds to the scalar density values at each voxel, with the color intensity indicating density magnitude.

In ***Figure 5*Figure 5**, CSFSC curves are plotted to assess the resolution of the obtained 3D density maps. Different variations of Fourier Shell Correlation (FSC) plots are presented: one employing an automatically generated mask with a 15 Å falloff, termed the ‘loose mask’ curve, and the other using an auto-generated mask with a falloff of 6 Å for all FSC plots, referred to as the ‘tight mask’ curve. The 3D density map reconstructed by each method for each protein is also visualized. The notable difference between the results of CrYOLO and CryoTransformer can be observed. For instance, in the case of EMPIAR 10345, the correct shape of the density map has three distinct legs, but CrYOLO failed to capture all three, yielding a lower resolution of 6.06 Å. In contrast, CryoTransformer captured all of them and achieved a high resolution of 3.45 Å. Similarly, in case of EMPIAR 10532, Topaz missed the central segment of the rod-like protein structure, whereas CryoTransformer successfully reconstructed that portion, attaining the highest resolution (3.21 Å) among all methods.

The plot located in the lower-right corner of each section in **Figure 5** represents the intermediate output of the ab-initio reconstruction phase. These plots depict density projections, but instead of slicing the density along a specific plane, the integrated density values along the normal direction to that plane are displayed. The color scheme in the heatmap corresponds to the scalar density values at each voxel, with color intensity indicating density magnitude.

In addition to this visual assessment in **Figure 5**, we conducted a comparison based on the 2D orientation classes of the picked particles (**Figure 6)**, showing that CryoTransformer picked particles in multiple orientation states that are important for obtaining high-resolution density maps. This analysis specifically involved analyzing the elevation vs azimuth plots for each test EMPIAR IDs. In the case of EMPIAR 10532 row, CrYOLO struggled to select particles representing various orientations, resulting in low-quality 2D particle classes. In contrast, Topaz performed reasonably well in picking particles with a diverse range of orientations, and CryoTransformer excelled in picking a substantial number of particles with a broad angular distribution, as indicated by the red color in the heatmap. The higher intensity of the red color in the upper section of each blocks in **Figure 6** corresponds to the higher number of particles in the elevation vs azimuth plots. Similarly, the lower section of each block depicts the averaged 2D orientation classes generated from picked particles. The diverse set of particles picked by CryoTransformer enabled the reconstruction of the density map of the highest resolution for this protein.

**Figure 6:**
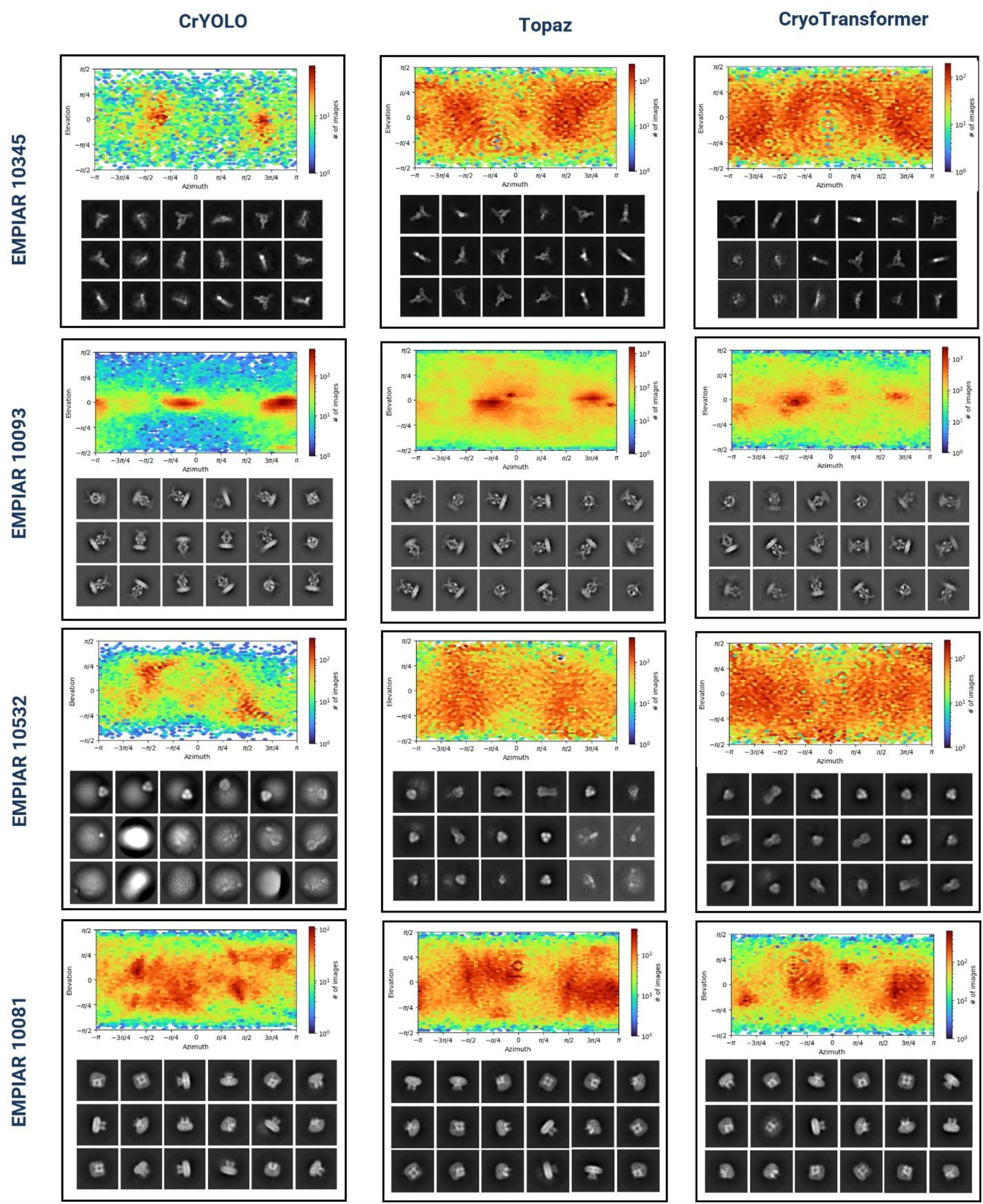
Assessment of CrYOLO, Topaz, and CryoTransformer based on the 2Dl orientation classes of the picked protein particles. Each block displays two sections: the upper section presents the viewing direction plots as elevation vs azimuth plots, while the lower section showcases the averaged 2D orientation classes generated from picked particles.

### 3.2. Comparing CryoTransformer, CrYOLO, and Topaz on a subset of micrographs in CryoPPP for the independent test proteins (~300 micrographs per protein)

Similarly, as in Section 3.1, we compared CryoTransformer, CrYOLO, and Topaz on the labeled subset of micrographs in CryoPPP for the six proteins in the independent test dataset in terms of the resolution of reconstructed density maps. The density maps were reconstructed using the *Select 2D* job from the picked particles. The 3D resolution is listed in **Table 3**.

**Table 3:** Comparison of CryoTransformer with CrYOLO and Topaz’s performance in terms of the resolution of 3D density maps reconstructed for six test proteins from the particles picked from a small set of micrographs in the CryoPPP.

| EMPIAR ID | Number of Micrographs | Number of Particles |  |  | 3D Resolution (Å) |  |  |
| --- | --- | --- | --- | --- | --- | --- | --- |
|  |  | CrYOLO | Topaz | CryoTransformer | CrYOLO | Topaz | CryoTransformer |
| 10017 [59] | 84 | 283 | 98,625 | 43,735 | 10.4 | **5.3** | 5.61 |
| 10081 [55] | 300 | 17,550 | 111,752 | 88,632 | 9.78 | 7.81 | **5.47** |
| 10093 [57] | 295 | 8,802 | 257,490 | 151,545 | 8.8 | **6.06** | 6.86 |
| 10345 [58] | 295 | 4,095 | 93,699 | 105,739 | 10.2 | 8.12 | **6.43** |
| 10532 [56] | 300 | 12,166 | 356,222 | 148,345 | 15.69 | 5.47 | **3.92** |
| 11056 [60] | 305 | 46,582 | 144,600 | 98,193 | 10 | 8.34 | **7.42** |

Among the six datasets considered, CryoTransformer outperforms crYOLO and Topaz in four instances, despite picking a much smaller number of particles than Topaz in most cases. This observation underscores Topaz’s tendency to pick more overlapped/duplicate particles or false positives. CrYOLO performs substantially worse than CryoTransformer and Topaz because it picks a much small number of particles, which are not sufficient to build good density maps. For the same four proteins, the best resolution of the density maps in **Table 3** is lower than that in **Table 2** because a much small number of micrographs were used for the particle picking and density map reconstruction.

In addition to the evaluation based on 3D resolution and the number of picked particles, we also assessed the performance of the three methods using precision, recall, F1 score, and dice score, as detailed in **Table 4**.

**Table 4:** Comparison of CryoTransformer with crYOLO and Topaz in terms of precision, recall, F1-score, and dice score of particle picking on the micrographs of six independent test proteins.

| EMPIAR ID | Type of Protein | Number of Micrographs | # Particles in Ground Truth (CryoPPP dataset) | Precision |  |  | Recall |  |  | F1 Score |  |  | Dice Score |  |  |
| --- | --- | --- | --- | --- | --- | --- | --- | --- | --- | --- | --- | --- | --- | --- | --- |
|  |  |  |  | CrYOLO | Topaz | CryoTransformer | CrYOLO | Topaz | CryoTransformer | CrYOLO | Topaz | CryoTransformer | CrYOLO | Topaz | CryoTransformer |
| 10017 | β-galactosidase | 84 | 49,391 | 0.695 | 0.57 | **0.745** | 0.024 | **0.998** | 0.587 | 0.046 | **0.726** | 0.657 | 0.041 | **0.694** | 0.623 |
| 10081 | Transport | 300 | 39,352 | 0.405 | 0.736 | **0.860** | 0.18 | **0.965** | 0.889 | 0.25 | 0.835 | **0.874** | 0.214 | **0.825** | 0.823 |
| 10093 | Membrane | 295 | 56,394 | 0.574 | **0.617** | 0.560 | 0.054 | **0.537** | 0.689 | 0.098 | 0.574 | **0.618** | 0.086 | 0.504 | **0.600** |
| 10345 | Signaling | 295 | 15,894 | 0.543 | 0.526 | **0.744** | 0.134 | **0.981** | 0.864 | 0.215 | 0.685 | **0.799** | 0.111 | 0.659 | **0.684** |
| 10532 | Viral | 300 | 87,933 | 0.715 | 0.672 | **0.813** | 0.201 | **0.976** | 0.665 | 0.313 | **0.796** | 0.732 | 0.239 | **0.757** | 0.614 |
| 11056 | Transport | 305 | 125,908 | 0.513 | 0.731 | **0.853** | 0.214 | **0.832** | 0.683 | 0.302 | **0.778** | 0.758 | 0.284 | **0.692** | 0.679 |
|  | Average |  |  | 0.574 | 0.642 | **0.7625** | 0.1345 | **0.8815** | 0.7295 | 0.204 | 0.732 | **0.740** | 0.163 | **0.689** | 0.671 |

Moreover, we compared the particles picked by each method with the ground truth particles labeled in CryoPPP in terms of four machine learning metrics including precision, recall, F1 score (the geometric mean of the precision and recall), and Dice score (**Table 4**).

CryoTransformer stands out with the highest average precision of 0.7625 and the highest average F1-score of 0.740, indicating that it excels in producing accurate positive predictions and achieves the best-balanced performance considering both precision and recall. Topaz has the highest average recall and dice score of 0.8815 and 0.671, highlighting its ability to correctly identify a high proportion of true positive particles and generate a strong overlap between predicted and actual positive instances.

### 3.3 Visual inspection of particles picked by CryoTransformer, CrYOLO, and Topaz

Figure 7. visualizes the particles picked by the three methods from the four representative micrographs of four proteins in the internal test data, which consist of 10% of micrographs from the 80%-10%-10% train-valid-test split (see detailed results in **Supplementary Table S4**). Consistent with the results in **Section 3.2**, CrYOLO tends to select few true protein particles, consequently missing many true positive across various protein types. In contrast, Topaz selects an excessive number of particles, often leading to overlaps and duplicates. Additionally, Topaz frequently picks false particles from contaminations, particle aggregates and ice patches, which can result in lower-resolution 3D density map reconstruction. Furthermore, picking a lot of redundant particles requires much more storage to store them and a lot of time and memory to reconstruct density maps from them. CryoTransformer, on the other hand, often picks most of true particles while keeping false positives to a low level, which is beneficial for 3D density map reconstruction.

**Figure 7:**
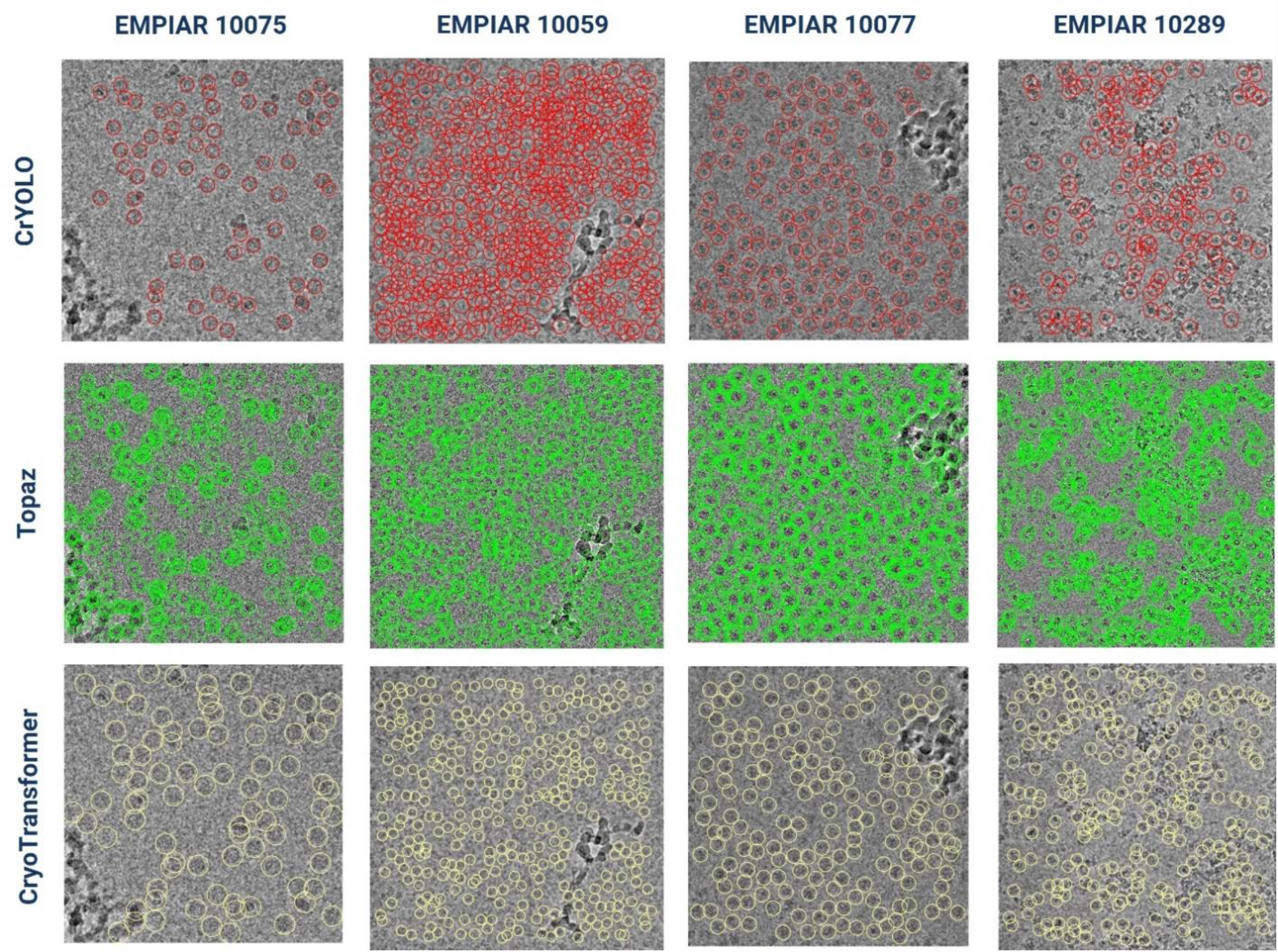
Assessment of CrYOLO, Topaz, and CryoTransformer based on visual inspection of predicted particles in micrographs of four typical proteins. The first row (indicated by red circles) represents protein particles picked by CrYOLO. The second row (marked by green circles) displays protein particles picked by Topaz. The third row (with yellow circles) illustrates protein particles picked by CryoTransformer.

## Conclusion

We present CryoTransformer, a novel deep learning method for particle recognition and extraction. It leverages the power of transformers, residual networks, traditional image processing techniques, and a bipartite matching loss function. CryoTransformer was trained and tested on the largest labeled particle dataset available. According to the rigorous evaluations and comparisons, CryoTransformer achieves state-of-the-art performance, making it a robust AI-based tool to automate the process of picking protein particles from cryo-EM micrographs.

## Code Availability

The source code and data are available at the GitHub repository: https://github.com/jianlin-cheng/CryoTransformer.

## Acknowledgements

We thank the EMPIAR team for maintaining the Cryo-EM data archive and the scientists who contributed their cryo-EM micrographs to EMPIAR repository. We are grateful to Dr. Filiz Bunyak for her insights on handling complex microscopic images through computer vision perspectives.

## Author Contributions

J.C. conceived and conceptualized this research; J.C. and L.W. provided guidance on the research and evaluation; A.D., R.G., and J.C. designed the methods; A.D. implemented the methods; A.D. performed the training and evaluations; A.D. and R.G. drafted the manuscript; J.C. and L.W. revised the manuscript; and all authors discussed the results and contributed to the final manuscript.

